# PACS1 syndrome variant alters proteomic landscape of developing cortical organoids

**DOI:** 10.1101/2025.04.23.650290

**Authors:** Ximena Gomez-Maqueo, Lauren E Rylaarsdam, Ashley Woo, Annika L Schroder, Jennifer Rakotomamonjy, Clare L Bossert, Tess A Smith, Shelby Ruiz, Jordan Gilardi, Lambertus Klei, Bernie Devlin, Matthew L MacDonald, Alicia Guemez-Gamboa

## Abstract

PACS1 syndrome is a neurodevelopmental disorder (NDD) resulting from a unique *de novo* p.R203W variant in Phosphofurin Acidic Cluster Sorting protein 1 (PACS1). PACS1 encodes a multifunctional sorting protein required for localizing furin to the *trans*-Golgi network. Although few studies have started to investigate the impact of the PACS1 p.R203W variant, the mechanisms by which the variant affects neurodevelopment are still poorly understood. In recent years, autism spectrum disorder (ASD) patient-derived brain organoids have been increasingly used to identify pathogenic mechanisms and possible therapeutic targets. While most of these studies evaluate the mechanisms by which ASD-risk genes affect the transcriptome, studies considering the proteome are limited. Here, we examine the effect of PACS1 p.R203W on the proteomic landscape of brain organoids using tandem mass tag (TMT) mass-spectrometry. Time series analysis between PACS1^(+/+)^ and PACS1^(+/R203W)^ organoids uncovered several proteins with dysregulated abundance or phosphorylation status, including known PACS1 interactors. Although we observed low overlap between proteins with altered expression and phosphorylation, the resulting dysregulated processes converged. The presence of the PACS1 p.R203W variant accelerated the emergence of proteins related to synaptogenesis and impaired vesicle loading and recycling. The earlier presence of these proteins and their related processes could lead to defective and/or incomplete synaptic function. Key dysregulated proteins observed in PACS1^(+/R203W)^ organoids have been associated with several neurological diseases, and many are classified as NDD-causative and ASD-risk genes. Our results highlight that proteomic analyses not only enhance our understanding of general NDD mechanisms by complementing transcriptomic studies, but could also uncover additional targets, and therefore facilitate therapy development.

## Introduction

Neurodevelopmental disorders (NDD) are heterogenous and usually present with complex etiology. Several large-scale genetic studies investigating the cause of NDDs such as autism spectrum disorder (ASD), intellectual disability (ID), and epilepsy have been conducted, and have lead to the identification of hundreds of risk genes. However, our understanding of the mechanisms by which the observed genetic variants act has lagged, limiting translation of genetic findings into clinical treatments (1, 2). Characterizing the pathogenic mechanisms of different monogenic NDDs may reveal convergent therapy targets. PACS1 syndrome (MIM615009) is a monogenic NDD characterized by ASD, ID, craniofacial abnormalities, and epilepsy (3–5). PACS1 syndrome results from a recurrent *de novo* c.607C>T variant in the gene encoding <u>P</u>hosphofurin <u>A</u>cidic <u>C</u>luster <u>S</u>orting protein 1 (PACS1). This missense variant is present in all reported patients and results in an arginine to tryptophan (p.R203W) substitution at residue 203, with the exception of one known arginine to glutamine p.R203Q substitution (5).

*PACS1* encodes a multifunctional sorting protein required for furin localization to the *trans*-Golgi network (tGN) (6, 7). PACS1 binds furin and other targets via acidic cluster sorting motifs (8–11). In addition to tGN retrieval, PACS1 mediates the recycling of endocytosed furin from early endosomes to the cell surface and the delivery of proteins to the primary cilium (12–14). PACS1 has been shown to interact with over 50 other proteins (10, 12, 15–17). Although the role of PACS1 has primarily been studied in the cytoplasm, several reports show that it also accumulates in the nucleus (18, 19) where it mediates DNA replication repair (20, 21), contributes to the nuclear export of viral transcripts (19), and regulates chromatin stability through interaction with histone deacetylase (HDAC) proteins (20).

Recent reports have begun to explore the pathogenesis of the PACS1 p.R203W variant. Mice overexpressing PACS1 p.R203W exclusively in cortical neurons present with increased dendrite arborization and decreased postsynaptic currents, possibly due to a pathogenic interaction between PACS1 p.R203W and HDAC6. Moreover, antisense oligonucleotides (ASO) directed towards both PACS1 and HDAC6 ameliorate these defects (22). Transcriptomic analysis in patient-derived brain organoids revealed a dysregulation in cell projection organization in immature PACS1 p.R203W neurons, while mature neurons exhibited an upregulation in synaptic signaling processes when compared to isogenic controls. Subsequent characterization of spontaneous electrical activity revealed that induced pluripotent stem cell (iPSC)-derived neurons harboring the p.R203W variant display asynchrony and an increased network burst duration (23). Finally, a study using *C. elegans* showed that the PACS1 syndrome-associated variant altered PACS1 localization but had no effect on neuron morphology and synaptic patterns (24). Although these studies have started to investigate the impact of the PACS1 p.R203W variant, the mechanisms by which the variant affects neurodevelopment are still poorly understood. This in turn results in few therapeutic options for patients, which at present, are limited to non-invasive behavioral approaches and sensory integration therapies to manage symptoms (25, 26).

In recent years, ASD patient-derived brain organoids have been increasingly used to identify pathogenic mechanisms and possible therapeutic targets (27). While most of these studies investigate the mechanisms by which ASD-risk genes affect the transcriptome (28, 29), studies considering the proteome are limited. However, mRNA levels do not always predict protein levels, as protein expression and degradation are regulated by mechanisms beyond transcription (30). Here, we uncover the proteomic landscape of brain organoids during maturation to provide further understanding into the pathogenic mechanisms of PACS1 syndrome.

## Methods

### Experimental Design and Statistical Rationale

The objective of this investigation is to provide further understanding of the pathogenic mechanisms of PACS1 syndrome by investigating proteomic landscape regulation during brain organoid maturation. We used tandem mass tag (TMT) mass-spectrometry in brain organoids generated from control and iPSC lines carrying the unique PACS1 p.R203W variant. Organoids were collected on days 21, 40, and 90 to better understand the maturation process. Protein and phosphorylated peptides were processed through mass-spectrometry. Outputs were normalized and aggregated by isogenic pairs in R using the package limma (31). Processed outputs were then fitted into a linear model and tested for differential expression over maturation time. Processed data were imported into the short time series (STEM) (32) pipeline for expression trend clustering (Figure S1).

### Generation of forebrain organoids from iPSCs

Three isogenic pairs of PACS1^(+/+)^ and PACS1^(+/R203W)^ iPSC lines were previously generated (23) and differentiated to brain organoids using the kit based on the protocol outlines by Sloan et al. (33) (Stem Cell Technologies, 08620). See Supplementary Methods for further details on experimental design, organoid generation, immunostaining and western blotting.

### Sample preparation for proteomic analysis

Total protein was extracted from the organoids by sonication in lysis buffer (1% sodium dodecyl sulfate (SDS), 125 mM triethylammonium bicarbonate (TEAB), 75 mM NaCl, and HaltTM protease and phosphatase inhibitors). Lysate samples from two additional control cell lines were included to improve protein detection. See Supplemental Methods and Supplementary Table 1 for a detailed description of mass spectrometry sample preparation, TMT labeling, and phosphopeptide enrichment processing.

### MS data acquisition and raw data processing

For peptide/protein quantification, TMT labeled peptides (∼1 μg) were loaded onto a heated PepMap RSLC C18 2 μm, 100 angstrom, 75 μm x 50 cm column (Thermo Scientific) and eluted over 180 min gradients optimized for each high pH reverse-phase fraction (34). Sample eluate were electrosprayed (2000 V) into a Thermo Scientific Orbitrap Eclipse mass spectrometer for analysis. MS1 spectra were acquired at a resolving power of 120,000. MS2 spectra were acquired in the Ion Trap with CID (35%) in centroid mode. Real-time search (max search time = 34 s; max missed cleavages = 1; Xcorr = 1; dCn = 0.1; ppm = 5) were used to select ions for synchronous precursor selection for MS3. MS3 spectra were acquired in the Orbitrap with HCD (60%) with an isolation window = 0.7 m/z and a resolving power of 60,000, and a max injection time of 400ms. For phosphopeptide quantification, 4 μl (out of 20) of the TMT labeled phosphopeptide enrichments were loaded onto a heated PepMap RSLC C18 2 μm, 100 angstrom, 75 μm x 50 cm column (Thermo Scientific) and eluted over a 180 min gradient: 1 min 2% B, 5 min 5% B, 160 min 25% B, 180 min 35% B. Sample eluate was electrosprayed (2000 V) into a Thermo Scientific Orbitrap Eclipse mass spectrometer for analysis. MS1 spectra were acquired at a resolving power of 120,000. MS2 spectra were acquired in the Orbitrap with HCD (38%) in centroid mode with an isolation window = 0.4 m/z, a resolving power of 60,000, and a max injection time of 350 ms.

The raw MS files were processed using Proteome Discoverer version 2.4 (Thermo Scientific, Waltham, MA). MS spectra were searched against the *Homo sapiens* Uniprot/SwissProt database. SEQUEST search engine was used (enzyme = trypsin, max. missed cleavage = 4, min. peptide length = 6, precursor tolerance = 10 ppm). Static modifications include carbamidomethyl (C, +57.021 Da) and TMT labeling (N-term and K, +304.207 Da for TMTpro16). Dynamic modifications include oxidation (M, +15.995 Da), Phosphorylation (S, T, Y,+79.966 Da, only for phosphopeptide dataset), acetylation (N-term,+ 42.011 Da), Met-loss (N-term, –131.040 Da), and Met-loss + Acetyl (N-term, –89.030 Da). PSMs were filtered by the Percolator node (max Delta Cn = 0y .05, target FDR (strict) = 0.01, and target FDR (relaxed)=0.05). Proteins were identified with a minimum of 1 unique peptide and protein-level combined q values < 0.05. Reporter ion quantification was based on corrected S/N values with the following settings: integration tolerance = 20 ppm, method = most confident centroid, co-isolation threshold = 70, and SPS mass matches = 65.

### Bioinformatic processing

Protein intensity data values were first normalized to adjust each TMT experiment to an equal signal per channel. A global scaling value and the internal reference scaling factor were then calculated (35), after which, a median normalization was performed and then log2-transformed. Further analysis of the data indicated a strong batch effect associated with multiplex labeling, which we ameliorated with the function removeBatchEffect in the R package limma (v3.58.0) (31). The experimental variability associated with genotype, day, isogenic cell line, and differentiation variables were retained in the design matrix to test for differential expression across genotypes and maturation time. Only the proteins (Supplementary Table 2-4) and phosphopeptides (Supplementary Table 5-7) present in all plexes were used for downstream analysis. This stringent filter was implemented to better control batch effects and addressed the impossibility of differentiating the missing values from non-expressed proteins. Despite this stringent filter, we captured a diverse pool of proteins and phosphopeptides similar to other publicly available datasets (27).

#### Time series clustering

We used the Short Time-series Expression Miner software (STEM, v1.3.13;) (32), which compares and assigns each expression profile from the dataset to a set of distinct and representative models of temporal expression profiles based on the total timepoints. These model profiles are independent of the data and correspond to possible profiles of a protein change in expression over time. For its implementation, the batch-corrected normalized protein abundances in the organoids were first log2-transformed and averaged by isogenic line and timepoint. We also used a fetal brain proteome previously reported by Djuric *et al*. (36) and selected the prenatal samples from all reported brain regions (ventricular zone, intermediate zone, subplate zone, and cortex). Fetal brain samples were processed using each region as a replicate for clustering. The STEM algorithm then transforms all trends to start at 0, generating a vector (0, *v_1_* — *v_0,_ v_2_* — *v_0_*, …, *v_n_* — *v_0_*) for each protein. Proteins are then assigned to the model profile to which their time series most closely matches based on correlation coefficient. The number of proteins assigned to each model profile is then tested for statistical significance against the number of proteins expected to be assigned to a profile. This number is estimated by randomly permuting the original time points, renormalizing the protein abundances, and then assigning them to the most closely matching model profiles. The average number of proteins assigned to a model profile over all permutations was used as the estimate of the expected number of genes assigned to the profile. This process was performed using a significance level of 0.05 of the Bonferroni-corrected p-value. The values used for each parameter are presented in Supplementary Table 8. Several preliminary runs were performed to select the values that both offered clustering of most proteins and diversity of profiles, while also controlling the number of highly similar profiles. Protein overlap among analyzed datasets were performed using the packages GeneOverlap (v.1.40.0) (37) and UpSetR (v.1.4.0) (38).

#### Functional processes – Gene enrichment analysis

Testing for enrichment was performed with the online tool Metascape (v3.5.20240901) (39) using a hypergeometric test followed by the Benjamini-Hochberg *p*-value correction, selecting statistically significant terms at a cutoff value below -1.3 Log(q-value) (39), and then simplifying redundancy in the resulting lists using rrvgo in R (v1.16) (40). Enrichment tests were performed using the Gene Ontology (GO) biological processes (41, 42) and the gene-disease association (DisGeNET) (43) databases. Visualizations were performed using webr (v.0.1.5) (44) and ggplot2 functions (v3.5.1) (45). The GO-term network was performed in Metascape. In brief, the 20 top-score clusters (lowest p-values) were selected from each dysregulated expression trend and then used the Kappa similarity statistic above 0.3 to connect similar terms (39). The resulting network was then edited in Cytoscape (v3.10.2) (46). Additionally, we consulted the SFARI Gene database (2024 Q3 release) (47) and OMIM (48) to detect dysregulated proteins that have been annotated as ASD-risk genes and/or pathogenic variants on such genes cause NDD.

## Results

### Brain organoids capture proteomic developmental trajectories and processes observed in fetal brain

Proteins are the ultimate effectors of cellular processes. Thus, here we characterized the proteome of iPSC-derived brain organoids to better understand how p.R203W PACS1 impacts neurodevelopment. Organoids were collected at distinct differentiation time points: day 21, when proliferating neural precursors predominate; day 40, when some progenitors have transitioned into intermediate progenitors and newly born neurons; and day 90, when post-mitotic neurons are the most abundant population (Fig. 1A).

**Figure 1.**
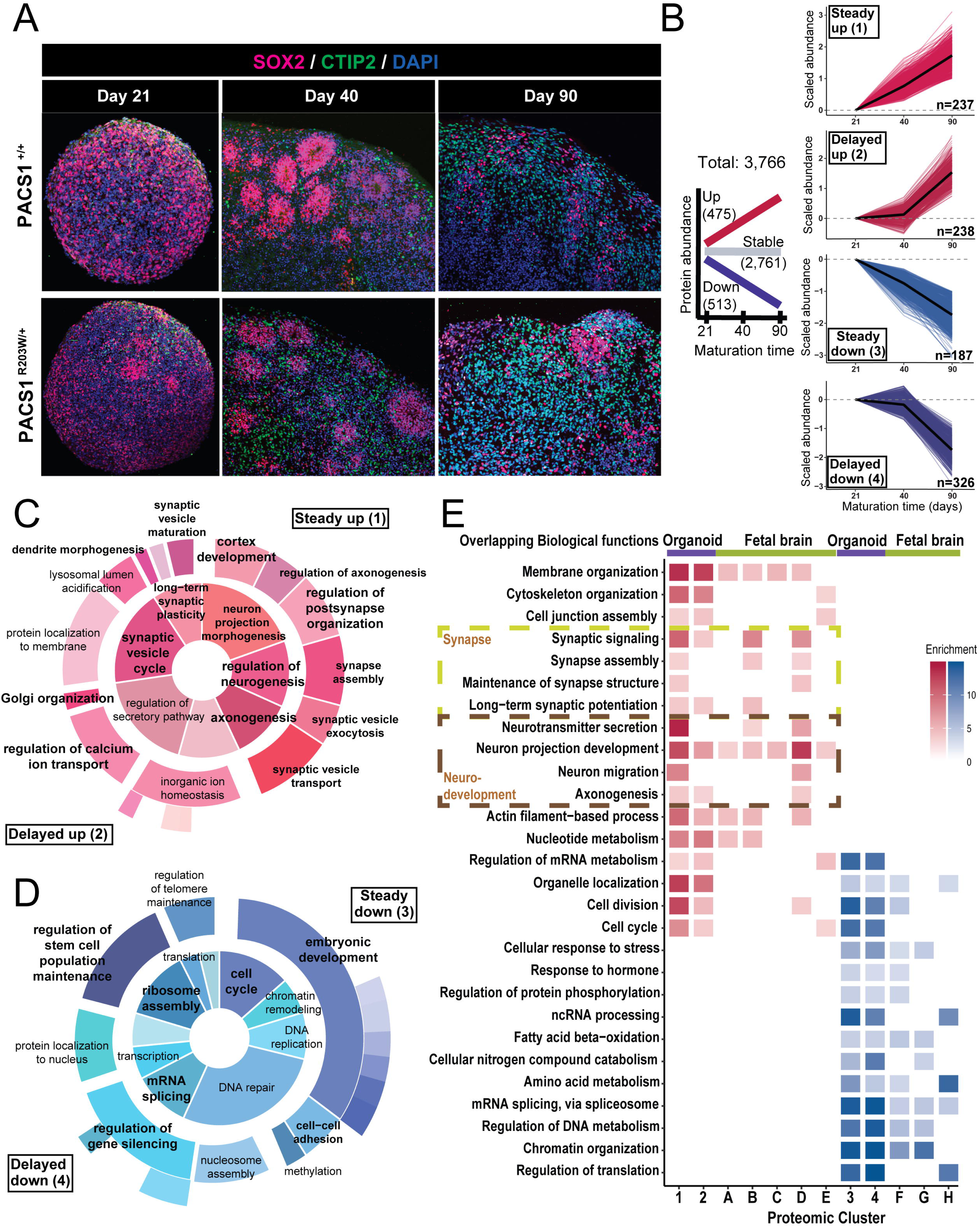
Cortical organoids recapitulate proteome dynamics during brain development. A) Expression of neural precursor protein SOX2 (red) and projection neuron marker CTIP2 (magenta) in day 21, 40, and 80 organoids derived from control (up) and PACS1 p.R203W organoids (bottom). Scale bar = 100 µm. DAPI (blue) B) Total proteins detected in organoids were clustered by abundance changes over maturation time into three major trends: upregulated (red), downregulated (blue) or stable (gray) (graph on the left). Further classification detected four main abundance trends, based on the timing of change: steady over time (clusters steady -up and -down), or delayed to day 40 (clusters delayed -up and -down) (Right). Total number of proteins assigned to a cluster are indicated in each graph. C, D) Top GO terms upregulated (red) and downregulated (blue) by cluster over maturation time. The inner pie plots represent processes common to the clusters in each panel. The nested pie charts show root and branch significant GO enrichments with the width of the pie correlating to -log(p) of the GO term. Processes in bold represent those mentioned in the text. E) Top GO terms upregulated (red) and downregulated (blue) shared between organoids (purple) and fetal brain (green) over time. Synapse and neurodevelopment-related processes (box). Organoid profiles correspond to clusters 1 to 4, while the fetal brain profiles correspond to clusters A to H. A full description of all enriched processes is included in Supplementary Table 10.

We utilized TMT mass-spectrometry to detect 3,766 unique proteins in our organoids across stages, with ∼60% overlapping proteins present in fetal brains (Fig. S2A). Most overlap was observed within organoids and brain samples from post-conception week 16 through 26, with protein abundance correlation increasing with organoid maturation (Fig. S2B). First, we aimed to understand the proteomic landscape of maturing brain organoids derived from neurotypical iPSCs. We identified groups of proteins with common temporal trajectories by performing time series and clustering analysis. We classified proteins into three temporal patterns: upregulation, downregulation, and stable abundance over time. The up and downregulated patterns were further clustered into two main profiles: 1) continuous protein change, with steady increase/decrease in abundance (clusters “steady-up” and “steady-down”); and 2) delayed trend, where protein abundance remained stable during early differentiation (from day 21 to day 40), and subsequently increased/decreased at day 90 (clusters “delayed-up” and “delayed-down”) (Fig. 1B; Supplementary Table 9).

Next, we used gene ontology (GO) analysis to determine the biological processes involved within these distinct patterns (Supplementary Table 10). As expected, many developmental processes changed with time, including neuron differentiation, axon extension, dendrite morphogenesis, synaptic organization/function upregulation (Fig. 1C), and cell cycle and stem cell population maintenance downregulation (Fig. 1D). This is consistent with the differentiation from neural precursors to neurons during organoid maturation. Within the upregulated clusters, the steady-up contained most of the neurodevelopmental processes, while the delayed-up was mostly associated with signaling. Likewise, clusters steady-down and delayed-down shared a downregulation of cell cycle and DNA replication or repair (Fig.1D). Stable proteins were associated with basic functions like primary metabolism, regulation of gene expression, and ribosome/translation-related processes (Fig. S2C).

Finally, we compared the observed organoid processes with the developing fetal brain dynamics obtained from a reference dataset (36) (Fig. S2D and Supplementary Table 11). We found significant overlap between clusters in organoids and the fetal brain (Supplementary Table 12). The majority of enriched processes during fetal brain development were captured in the corresponding organoid clusters. These results confirm that our brain organoids recapitulate early stages of brain development, as previously reported (27), supporting their use to model protein dysregulation in PACS1 syndrome.

### PACS1 p.R203W pathogenic variant putatively accelerates neural maturation

We then profiled the proteome of PACS1 syndrome patient-derived brain organoids to evaluate the effects of the PACS1 p.R203W pathogenic variant on protein regulation during development (Fig. 1A). Organoids carrying the p.R203W variant developed similarly to control organoids by day 21, with only five proteins (HMGB3, HTATSF1, HM13, TCEAL3 and TCEAL5) showing upregulation (Fig. S3A). However, upon maturation, 329 proteins in PACS1^(+/R203W)^ organoids showed dysregulation when compared to controls (Supplementary Table 13). Oxidative phosphorylation (ATP6V0A1, CYC1, NDUFA7) and intracellular calcium ion homeostasis (ATP1B1, CALB2, LETM1, NPTN) were among the observed enriched processes resulting from 167 upregulated proteins in organoids carrying the p.R203W (Fig S3B, Supplementary Table 14). By day 40, most dysregulated proteins were related to established PACS1 functions relating to endoplasmic reticulum (ER)-to-Golgi transport (ARF4, SEC23A, SNAPIN). Upregulated proteins at day 40 were involved in neuron projection development, such as the ligand of Eph-related receptor tyrosine kinases EFNB1 and the transcription factor FOXG1. This was confirmed to be upregulated by western blot (Fig S3C). In contrast, neuron projection development and synapse signaling related proteins like EFNB3, NF1, VDAC3, and SV2A were enriched by day 90. Other upregulated processes include the metabolism of amino acids (ASS1, GLO1, CTH, HSD17B10, PLOD2, SMS), and glycerolipids (PITPNM1, AGPAT4, TMEM68). Interestingly, the PACS1 interactors PTBP1 (49), involved in mRNA processing, and WDR37 (50), were also upregulated (Fig S3A,B). On the other hand, 173 down-regulated proteins were enriched in processes including cell cycle division, DNA replication, and mRNA splicing. This further included the RNA-binding protein PCBP4, involved in mRNA stability (51) and histone deacetylase PTBP2, which is also involved in brain development. The intermediate filament protein NES, which plays a role in proliferation and differentiation (52), was also found to be among the downregulated proteins. Decreased abundance of NES was confirmed by western blot (Fig S3).

Principal component analysis (PCA) indicated maturation time to be the strongest driver of variance when comparing PACS1^(+/+)^ and PACS1^(+/R203W)^ organoids at days 21, 40, and 90 (42% of variation) (Fig. S4A). Consequently, we investigated whether the presence of the PACS1 variant impacts groups of proteins that share common temporal trajectories. PACS1^(+/R203W)^ organoids also presented with the three major patterns (upregulation, downregulation, and stable) and profiles (steady vs. delayed) observed in controls (Fig. 2A, Supplementary Table 15). However, several proteins switched abundance patterns (Fig. 2B). The profile rearrangement resulted in four additional clusters, for a total of eight clusters found in PACS1^(+/R203W)^ organoids. Additional clusters included proteins that were up or down-regulated when compared to the stable pattern in controls (“stable-up” and “stable-down” clusters) and those that lost the regulation observed in controls and remained stable over maturation (“up-stable” and “down-stable” clusters; Supplementary Table 16).

**Figure 2.**
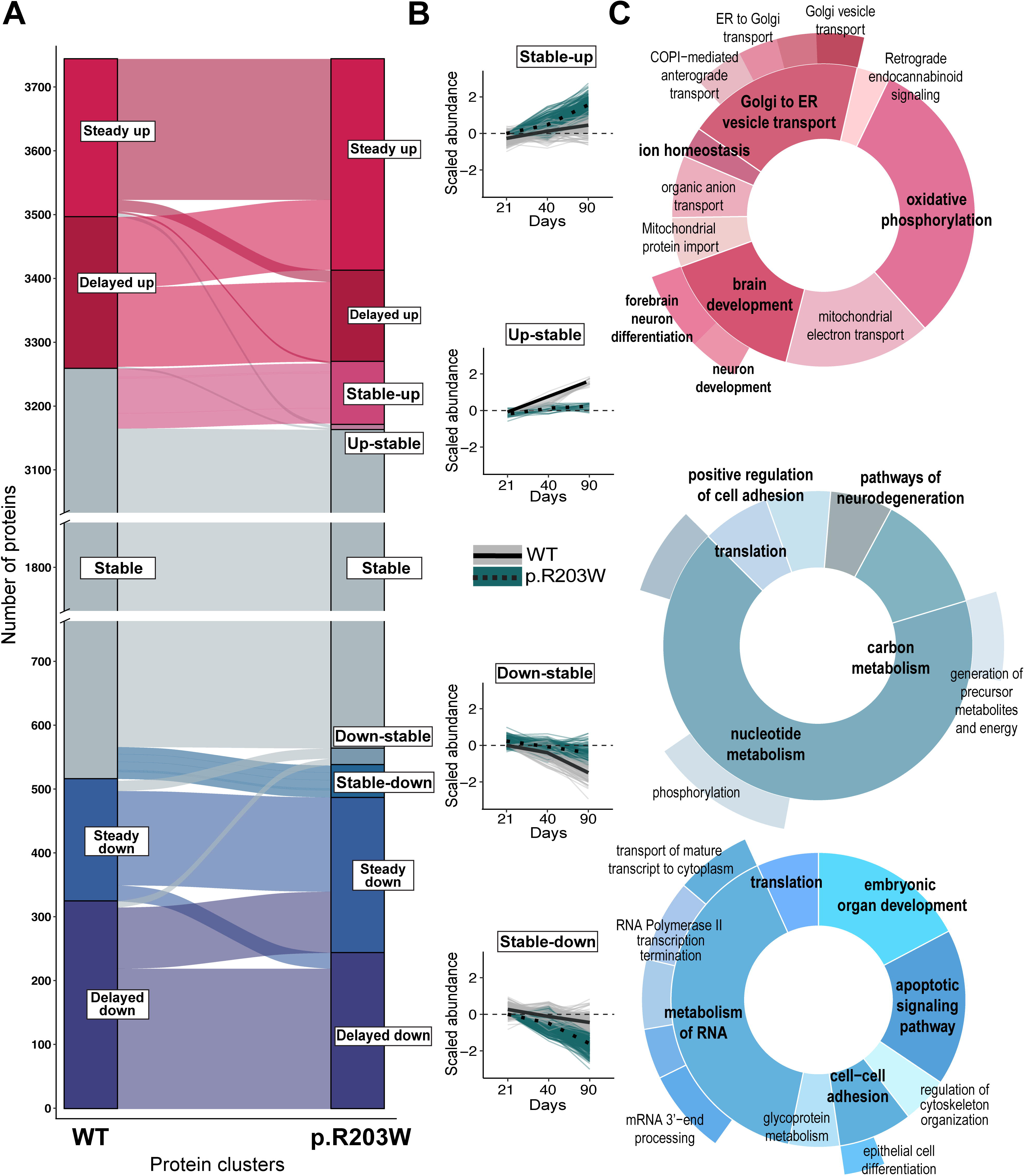
Organoids carrying the PACS1 p.R203W pathogenic variant present with altered protein dynamics during maturation. A) Alluvial plot indicating the change of profile clustering in the PACS1 p.R203W organoids (right) in contrast to the clustering in controls (left). Breaks in the y axis were introduced to improve visualization of the dysregulated clusters by removing 2,000 proteins that remained stable over time. B) Four clusters of dysregulated proteins were identified in PACS1 p.R203W organoids. The plots are organized into a gain or loss of regulation categories: cluster stable-up (top), and stable-down (bottom) gained regulation while clusters up- and down-stable lost regulation (middle), C) Enriched biological functions per dysregulated cluster Stable-up (top, red), Down-stable (middle, greyish blue) and Stable-down (bottom, blue). The Up-stable cluster had no significant enrichment due to cluster size. The nested pie charts show root and branch significant GO enrichments and the width of the pie correlating to -log(p) of the GO term. Neurodevelopment-related processes in (bold).

The stable-up cluster from PACS1^(+/R203W)^ organoids had a strong enrichment in oxidative phosphorylation (23 out of 33 proteins), brain development, and neuron differentiation. Proteins that gain upregulation include glucose transporter GLUT1, transcription factor FOXG1, and mitochondrial genes UQCRQ, VDAC1, and ATP5PF (Fig. 2C, Supplementary Table 17). The stable-down cluster was enriched in regulation of mRNA processing and translation. Proteins that gain downregulation include ribosomal subunit protein RPL38, alpha subunit of the elongation factor-1 complex EEF1A1, and developmental proteins Nestin (52), BRD2 (53), and Wnt family member 7B WNT7B (54).

The down-stable cluster was enriched in the metabolism of the nucleotides process. Proteins related to metabolism that lost their down regulation and remained stable over time have also been associated with brain development and function. These proteins include, ribokinase (RBKS), implicated in glucose metabolic disorders and recently associated to psychiatric disorders (55), Phosphoribosylpyrophosphate synthetase 1 (PRPS1), implicated in multiple syndromic and non-syndromic diseases causing hearing loss, hypotonia, and ataxia (56), as well as catalase (CAT), implicated in oxidative stress-related diseases and ASD (57).

Finally, the up-stable cluster was comprised of eight proteins which lost their temporal upregulation and remained stable in PACS1^(+/R203W)^ organoids, with no significant enrichment. In this group, however, several proteins have relevant functions in the brain, such as dendrite morphogenesis (SS18L1/CREST) (58), autophagy and neuron projection regeneration after injury (VIM) (59), and regulation of intracellular pH homeostasis (CA2) (60).

Our results uncovered the dysregulation of known PACS1 interactors at the protein level that were not resolved on the transcriptome level (23). Results suggest that the PACS1^(+/R203W)^ organoids undergo accelerated development, with the upregulation of processes related to known PACS1 functions (i.e., Golgi to ER vesicle transport). However, even if the PACS1^(+/R203W)^ organoids present with early neural maturation signatures, they also lost regulation of many mRNA related processes, like transport and processing, which could lead to defective and/or incomplete differentiation (Supplementary Table 17).

### Temporally regulated phosphorylation of proteins related to axonal and synaptic processes are also impacted in PACS1^(+/R203W)^ brain organoids

Post-translational phosphorylation is a main regulator of protein function. Thus, we analyzed whether the PACS1^(+/R203W)^ organoids also presented with alterations in phosphorylation during maturation. Quantification of phospho-peptides in organoids resolved a total of 4,238 phosphorylation sites (Supplementary Table 5). The time series and clustering analysis uncovered the same eight clusters and stable patterns that resulted from the protein abundance analysis (Fig. 3A, Supplementary Tables 18-20). However, an additional distinct cluster composed of peptides that acquired phosphorylated sites between days 21 and 40 (“early-up” cluster) was observed in PACS1^(+/R203W)^ organoids (Fig. 3B).

**Figure 3.**
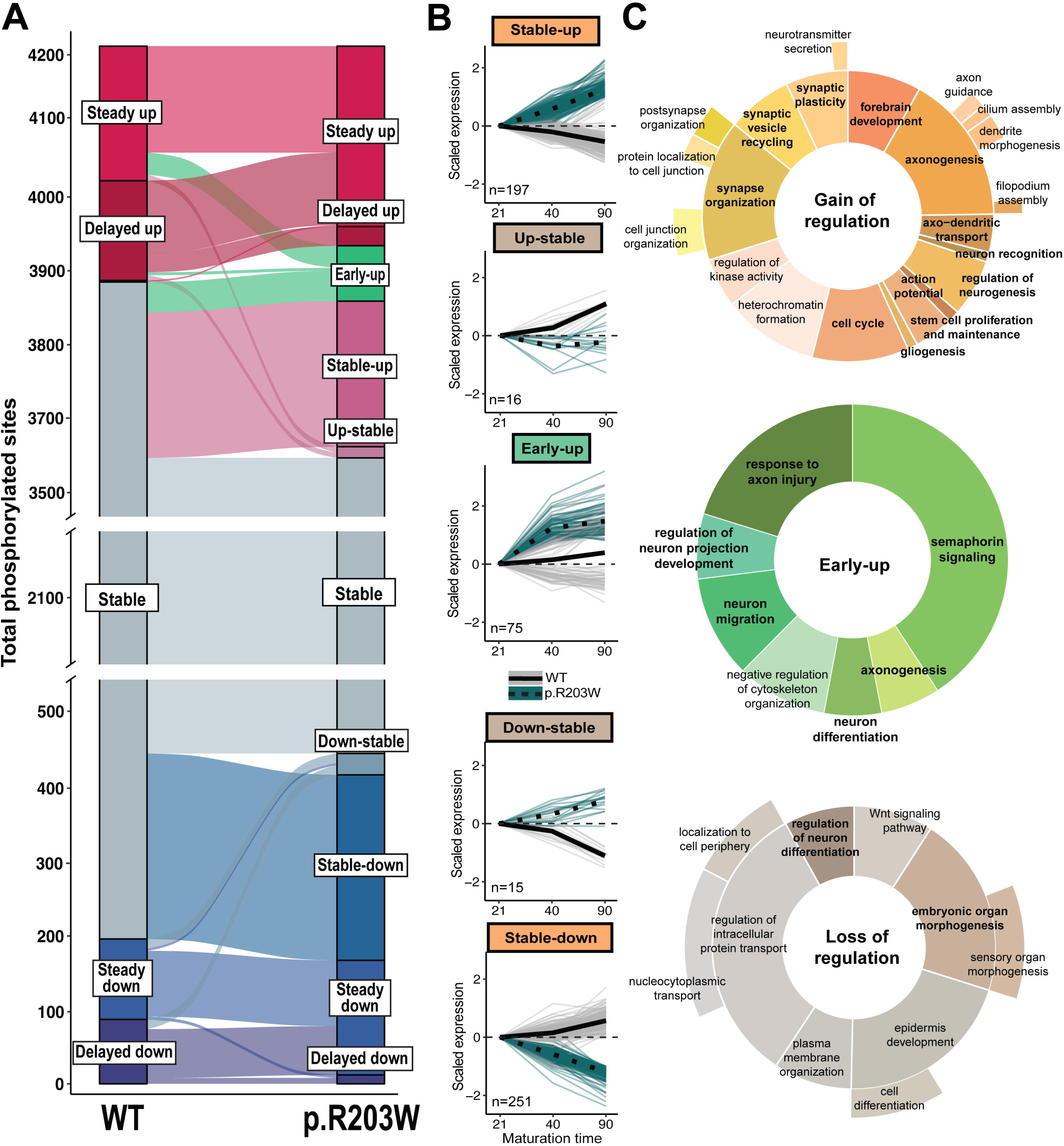
Dysregulation of temporal phosphorylation dynamics in PACS1 p.R203W organoids. A) Alluvial plot indicating the change of profile clustering in the PACS1 p.R203W organoids in contrast to controls. Breaks in the y axis were introduced to improve visualization of the dysregulated clusters by removing 3,000 modifications that remained stable over time. B) Five clusters of dysregulated phosphorylation trends were observed in PACS1 p.R203W organoids that either gained a temporal regulation or lost the original pattern observed in the controls. The plots are organized into a gain or loss of regulation categories: clusters stable- up and -down gained temporal regulation in contrast to the controls (orange), while clusters up- and down-stable lost temporal regulation (gray). Early-up is a new cluster exclusive to the phosphorylation dataset that also gained temporal regulation (green). C) Enriched biological functions per dysregulated cluster. The nested pie charts show root and branch significant GO enrichments and the width of the pie correlating to -log(p) of the GO term. Neurodevelopment-related processes (bold).

A total of 530 phosphorylation sites on 325 unique proteins were dysregulated in PACS1^(+/R203W)^ organoids (Supplementary table 20). Examples include two PACS1-known interactors, PACS2 (PACS1 paralog) and HDAC1. PACS1^(+/R203W)^ organoids presented with increased PACS2-S437 phosphorylation, which is required for PACS2-mediated trafficking and repression of apoptosis (61). HDAC1 had decreased phosphorylation at residues S421 and S423, which is essential for its repressive function (62). Interestingly, both serine residues are also phosphorylated by CK2 (found to have lost its temporal regulation and remained stable in PACS1^(+/R203W)^ organoids).

Because phosphorylation could result either in an induction or inhibition of protein function, and the activation status of a protein itself could have diverse effects on pathway activity, we used the Phosphosite database to understand the functional impact of the observed dysregulated phosphorylation sites (63). Phosphosite contained annotations on 51 out of 530 phosphorylation sites across 41 unique proteins, with few proteins having multiple phosphorylated sites (Fig. S5, Supplementary Table 21). To perform the functional enrichment analysis, we grouped phosphorylated proteins that gained (clusters: stable -up and -down) or lost (clusters up- and down-stable) phosphorylation regulation in contrast to the controls. Although the early-up cluster also gained upregulation, due to its uniqueness to this dataset, we performed the functional enrichment analysis separately (Fig. 3C, Supplementary Table 22).

The phosphorylated proteins that gained up/down-regulation in the PACS1^(+/R203W)^ organoids over time while remaining stable in controls are involved in axonogenesis, as well as synaptic function and organization processes (Fig. 3C, orange pie plot). Examples of these proteins include synapsin1 (SYN1), implicated in both synaptogenesis and neurotransmitter release, whose phosphorylation at residues S568/S570 causes conformational changes that prevents its binding to actin or synaptic vesicles (64); synaptojanin1 (SYNJ1), which has a prominent role in synaptic vesicle recycling (65); the beta subunit of the AP-3 protein complex, a known PACS1 interactor that helps form clathrin-coated synaptic vesicles (65); and DNM1-S774, which regulates clathrin-mediated endocytosis through its activation by dephosphorylation (66).

Similarly, as observed in the other gain of regulation clusters, the early-up cluster had enrichment for axon-related function and organization, but showed novel terms related to signaling through semaphorin 3 (SEMA3E) (Fig. 3C, green pie plot). This result is in consensus with single-cell RNA sequencing analysis, which identified *SEMA3E* to be upregulated in PACS1^(+/R203W)^ brain organoids (23). Our proteomic dataset shows that while there are no changes in SEMA3E, several proteins involved in its signaling pathway are phosphorylated, including CRMP1, DPYSL2, and DPYSL3. DPYSL2-S522 phosphorylation by CDK5 is involved in axonal guidance and spine development (67).

Meanwhile, intracellular protein transport was the most enriched process in the clusters that lost regulation (Fig. 3C, brown pie plot). Proteins that remained stable in PACS1^(+/R203W)^ organoids include GLI3, which has a dual function as a transcriptional activator and a repressor of the sonic hedgehog (Shh) pathway dependent on phosphorylation (68) and LIN28A, an RNA-binding protein that is localized to the peri-endoplasmic reticulum and has been recently described as a global suppressor of genes in the secretory pathway (69).

### Dysregulated protein abundance and phosphorylation status in PACS1^(+/R203W)^ organoids result in convergent processes and overlap with known NDD-related genes

We next analyzed the overlap of protein identities with dysregulated abundance and phosphorylation status in PACS1^(+/R203W)^ organoids. Surprisingly, we observed a low overlap of both the total and dysregulated proteins identified in both datasets (Fig. 4A, B). Nevertheless, several affected processes converged (Fig. S6). For example, neurogenesis and axon-related processes are enriched due to both dysregulated protein abundance and phosphorylation status over time in PACS1^(+/R203W)^ organoids when compared to controls (Fig. S6 purple and yellow circles). Other converging processes include mRNA transport, actin and microtubule organization, and cell cycle dynamics. However, exceptions were also present, where processes will be representing mostly protein abundance or phosphorylation status. These exceptions included oxidative phosphorylation and energy production resulting from the dysregulation of protein abundance (purple); and chromatin remodeling and synapse organization, which are composed of proteins showing a gain of upregulation in phosphorylation status over time (yellow) (Fig. S6). These results suggest that PACS1^(+/R203W)^ organoid development is disrupted at different regulation levels.

**Figure 4.**
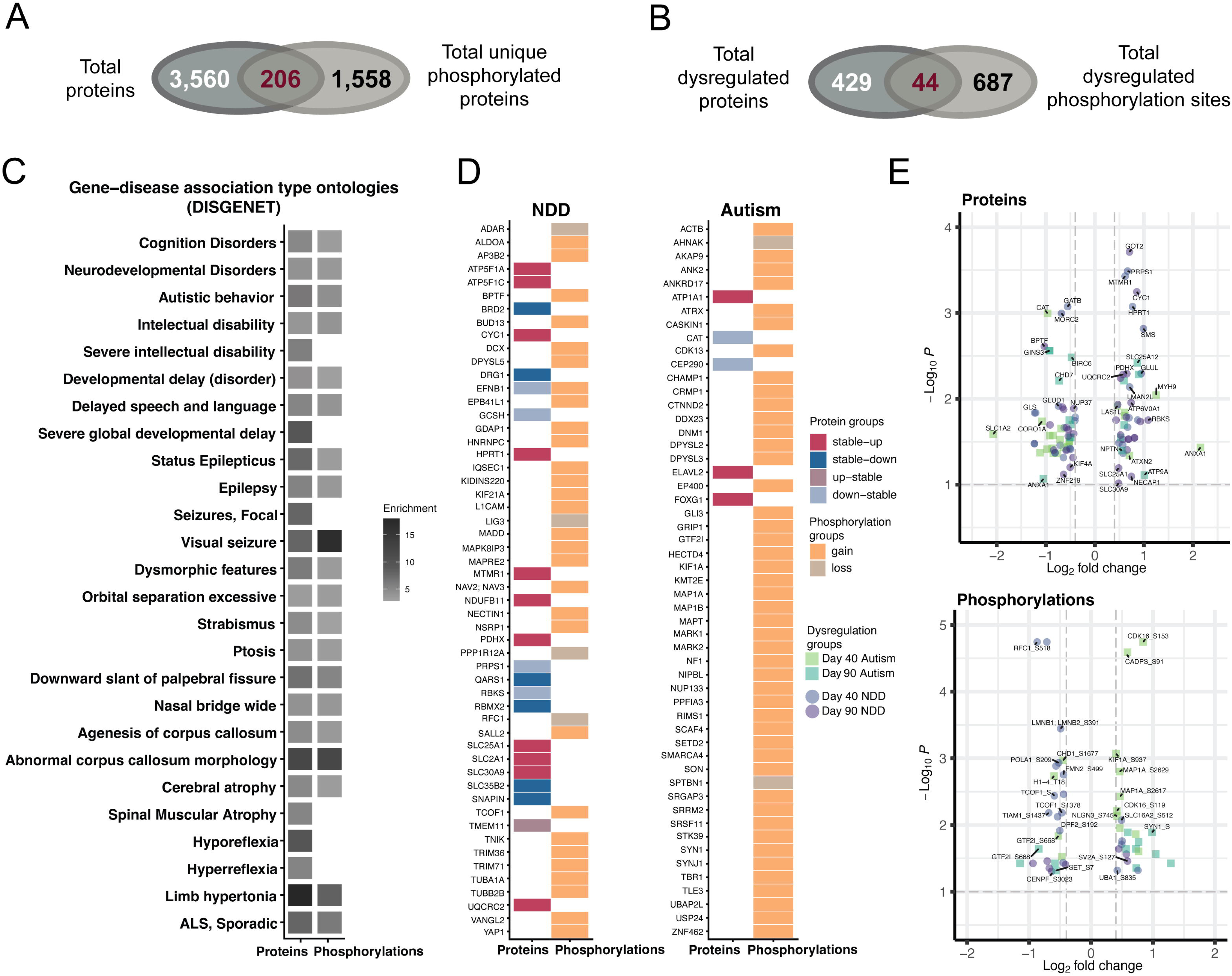
Protein abundance and phosphorylation sites dysregulated by PACS1 p.R203W are enriched in neurological disease-related genes. A) Overlapping identities between all detected proteins and proteins with identified phosphorylation sites. B) Overlapping identities between dysregulated proteins and phosphorylation sites observed in the PACS1 p.R203W organoids when compared to controls. C) Gene-disease association type ontologies enriched in the PACS1 p.R203W dysregulated proteome and phosphorylation sites. Selected top neuro-related categories at a corrected p-value of 0.01. D) NDD and ASD risk genes dysregulated in organoids carrying the p.R203W pathogenic variant. Genes are grouped by proteins and phosphorylations. The protein abundance group is colorized by dysregulation cluster while the phosphorylation clusters are colorized by major dysregulation trend (gain or loss) as presented in figure 3: gain of regulation (clusters stable- up and -down) or loss of regulation (clusters up- and down-stable). E) Volcano plot representing the dysregulated proteins (upper panel) and phosphorylations (lower panel) at days 40 and 90. The squares in green tones represent the ASD genes, while the circles in purple tones represent the NDD genes.

Next, we investigated whether there are convergent dysregulated proteins in PACS1 syndrome compared to other NDDs. We performed gene-set enrichment analysis on the dysregulated proteins using the DISGENET, the Simons Foundation ASD (SFARI), and the Online Mendelian Inheritance in Man (OMIM) gene databases. A total of 311 genes (26.8%) dysregulated in the PACS1^(+/R203W)^ organoids compared to controls are additionally reported in these databases. In DISGENET, we found 212 genes that showed a significant enrichment in multiple nervous system disorders (Fig. 4C; Supplementary Table 23). These disorders overlapped with phenotypes observed in PACS1 syndrome patients like ASD, ID, seizures, and abnormal morphogenesis of the nervous system or facial features (4, 70), as well as phenotypes yet to be reported, such as neurodegeneration. Finally, 94 of the genes that we identified to be dysregulated in PACS1^(+/R203W)^ organoids are also reported as ASD-risk genes in the SFARI database, while 105 proteins had reported pathogenic NDD-related phenotypes in the OMIM database (Fig. 4D,E Supplementary Table 24). These results support the overlap of PACS1 syndrome and dysregulated neurodevelopmental processes with ASD and expand on phenotypes that patients could manifest later in life.

## Discussion

NDDs are highly heterogenous and can arise from variants across hundreds of genes (62, 63). Even so, several limitations have hampered our understanding of the mechanisms by which these variants act, limiting translation of genetic findings into the clinic (64, 65). Here, we leveraged the potential of brain organoids to model neurodevelopment and gain insights into the pathogenic mechanisms of PACS1 syndrome. By using mass spectrometry with TMT, we identified 7,230 unique proteins, of which 3,677 were quantified with 100% present call in our samples, a yield consistent with findings from other proteomic studies involving brain organoids (27, 66, 67). We further observed temporal trajectories that reflected the transitions from proliferating progenitor cells towards mature neuronal lineages, mimicking the temporal dynamics observed in the fetal brain.

PACS1^(+/R203W)^ organoids presented with dysregulated abundance and phosphorylation status of temporally regulated proteins, including PACS1 known interactors. Despite the low overlap between expression and phosphorylation alterations in PACS1^(+/R203W)^, most dysregulated processes converged. We observed upregulation of processes related to axonogenesis and synapse maturation, suggesting accelerated development in PACS1^(+/R203W)^ organoids. Although PACS1^(+/R203W)^ organoids presented with early synaptic formation signatures, the regulation of multiple synapse and vesicle-related proteins is affected. At the same time, many synaptic maturation processes are lost, possibly disrupting proper synaptic function. This is consistent with the observed phenotype in mice overexpressing PACS1 p.R203W (22); and the asynchrony and an increased network burst duration reported in neurons harboring the p.R203W derived from patient iPSC lines (23). Interestingly, the synaptic terms obtained from protein abundance and phosphorylation were also obtained from transcriptomic data, suggesting that the PACS1 p.R203W variant might affect multiple regulatory levels.

Proteomic approaches are usually bulk and lack single-cell resolution; our experimental set up also lacks cells that emerge later in development (astrocytes and oligodendrocytes) or arise from non-ectoderm layers (microglia). With this being the case, other model systems should be explored to determine how these cells contribute to PACS1 syndrome pathogenesis. Nevertheless, we identified multiple proteins that displayed temporal changes in abundance and phosphorylation status in developing PACS1^(+/R203W)^ brain organoids with respect to controls. Of further interest, most proteins that we found dysregulated in PACS1^(+/R203W)^ organoids have also been linked to NDDs (68–70). Additional mechanistic analyses are required to understand if these effects are direct or indirect.

Most proteins that gain upregulation over time, and their resulting dysregulated processes, could be linked to known PACS1 functions. For example, PACS1’s role in protein sorting explains the overrepresentation of vesicle-related processes in the stable-up cluster. Moreover, several established PACS1 interactors were uncovered as dysregulated proteins, such as its close paralog, PACS2. PACS1 and PACS2 evolutionarily emerged as a duplication in higher metazoans from an ancient PACS gene (66). PACS1 function primarily involves sorting from endosomes to the tGN, which showed temporal dysregulation of over 20 proteins in PACS1^(+/R203W)^ organoids. Meanwhile, PACS2 evolved cargo-sorting functions from the Golgi to ER and vesicular coat preference for COPII; both processes upregulated in PACS1^(+/R203W)^ organoids. The upregulation in these processes is consistent with the detected gain of PACS2 regulation through increased phosphorylation at S437, which favors PACS2 trafficking function (47). This increased phosphorylated PACS2 could explain the dysregulated vesicular transport observed in PACS1^(+/R203W)^ organoids. On the other hand, it could also explain the downregulation of apoptotic signaling and mitochondrial function (Figure 2C). Of interest, a recurrent *de novo* missense mutation in the regulatory middle region of PACS2 is associated with PACS2 Syndrome, another NDD (71). Further research aimed at understanding the interconnection of PACS1 and PACS2 function, as well as their dysregulation and compensatory effects in disease, will help shed light on the pathogenesis of both PACS1 and PACS2 syndromes.

The processes that lost downregulation over time were related to recently described PACS1 functions, including proteins related to the maintenance of genome stability and DNA repair via interaction with HDAC2, HDAC3, and HDAC6 (20, 22). Although we did not observe protein abundance and phosphorylation status changes in HDAC2, 3, or 6, we found downregulation in HDAC5 at day 40, and HDAC1 had decreased phosphorylation over maturation. This suggests PACS1 could also modulate DNA stability through these proteins. HDAC1 interacts with RB1 and SMARCA4 (both dysregulated at the phosphorylation level) in a repressive complex in resting neurons that is abolished in a calcium-dependent manner to enable transcription (72). Indicating that the PACS1 p.R203W variant could affect transcription regulation by destabilizing the repressive complex formed by HDAC1, RB1, and SMARCA4.

Processes unrelated to known PACS1 functions also emerged, such as the enrichment associated with regulating nucleotide metabolism and protein translation in the down-stable cluster. There is growing evidence of the impact of protein translation on the pathogenesis of neurodevelopmental disorders, including ASD and fragile X syndrome (73). However, this is the first time that either process has been implicated in the PACS1 syndrome pathogenesis. Moreover, PACS1^(+/R203W)^ organoids presented with changes in genes reported to also be dysregulated in other neurological diseases. The presence of the PACS1 p.R203W variant, at the proteomic level, seems to affect processes related to the clinical features that patients manifest, including ASD, ID, and seizures (3, 61), as well as phenotypes yet to be reported, such as neurodegeneration. Additional models are required to test mechanisms of PACS1 syndrome outside the cerebral cortex, like the observed hypoplasia of the cerebellum and the corpus callosum, or the dysmorphic facial features.

Processes that seem unrelated to PACS1 function, but which have displayed upregulation in the PACS1^(+/R203W)^ organoids, should be further explored. Considering current evidence suggests the PACS1 p.R203W pathogenic variant may result in a gain of function (22, 23, 74), there is increased interest in generating targeted therapeutic strategies aimed at suppressing its toxic effects (22, 25). Thus, processes that seem unrelated to PACS1 function but with observed upregulation in the PACS1^(+/R203W)^ organoids should be a point of interest. Understanding whether the proteomic dysregulation results from the toxicity associated with the variant or as a compensatory mechanism to maintain cellular homeostasis would be especially relevant. Finally, the neurological phenotypes associated with dysregulated protein abundance and phosphorylation in PACS1 p.R203W organoids are consistent with clinical characteristics observed in PACS1 syndrome patients. Additional downstream analyses that aim to understand the converging dysregulated processes could offer novel therapeutic strategies for PACS1 syndrome. Our results highlight that proteomic analyses not only enhance our understanding of general NDD mechanisms by complementing transcriptomic studies, but may also uncover additional targets, facilitating therapy development.

## Abbreviations

(ASO): Antisense oligonucleotide
(p.R203W): Arginine to Tryptophan substitution
(ASD): Autism spectrum disorder
(FBR): Furin binding region
(GO): Gene ontology
(DISGENET): Gene-disease association ontologies
(HDAC): Histone deacetylase
(iPSC): Induced pluripotent stem cell
(ID): Intellectual disability
(NDD): Neurodevelopmental disorder
(PACS1): Phosphofurin Acidic Cluster Sorting protein 1
(PWC): Post-conception week
(PCA): Principal component analysis
(STEM): Short Time-series Expression Miner
(SFARI): Simons Foundation ASD gene database
(scRNASeq): Single cell RNA Sequencing
(TMT mass spectrometry): tandem mass tag mass-spectrometry
(tGN): *trans*-Golgi network

## Acknowledgements

This research was supported in part through the computational resources and staff contributions provided for the Quest high performance computing facility at Northwestern University which is jointly supported by the Office of the Provost, the Office for Research, and Northwestern University Information Technology.

## Authors’ contributions

This work was conceptualized by AGG. XGM performed proteomics analysis with help from SR, BD, JG, and MLM, and generated the figures with the help of AW. LR cultured the organoids, collected samples, and prepared them for proteomic analysis. AW, ALS, JR, CLB, and TAS performed wet lab experiments (immunohistochemistry, western blotting, imaging) and data analysis. Writing was done by XGM and edited by AGG, MLM, LR, JR, ALS, TAS, and CLB. All authors read and approved the final manuscript.

## Availability of data and materials

Data is provided within the manuscript or supplementary information files and from the corresponding author on reasonable request.

## Supplemental Material

**Supplementary information.** Supplementary methods and supplementary figures 1-6 containing additional data.

**Supplementary tables.** Supplementary tables 1-24 contain the list of proteins and phosphorylation sites, as well as accompanying statistical analysis performed on the data.

