## Supplementary_methods_and_fig_1-6 for "PACS1 syndrome variant alters proteomic landscape of developing cortical organoids"

This file contains the following information:

- Supplementary methods
- Supplementary figures S1 – S6
- Supplementary References

**Supplementary methods**

***Organoid generation***

Three isogenic pairs of PACS1$^{(+/+)}$ and PACS1$^{(+/R203W)}$ iPSC lines were previously generated (1) and differentiated to brain organoids using the kit (Stem Cell Technologies, 08620). Briefly, an AggreWell™800 24-well plate (Stem Cell Technologies, 34811) was prepared and 3 million cells from each line were seeded per well. STEMdiff™ Forebrain Organoid Formation Medium was exchanged for 5 days and on day 6, embryoid bodies from each Aggrewell were filtered through a 37 µM strainer (Stem Cell Technologies, 27250) and transferred to two wells of a 6-Well Ultra-Low Adherent Plate (Corning, 3471) containing STEMdiff™ Forebrain Organoid Expansion Medium. STEMdiff™ Forebrain Organoid Expansion Medium was changed every other day until day 26 when it was replaced with STEMdiff™ Forebrain Organoid Differentiation Medium. Organoids were cultured in differentiation media until day 43. Following day 43, organoids were cultured in STEMdiff™ Forebrain Organoid Maintenance Medium until collection.

***Organoid cryosectioning and staining***

Organoids were fixed with 4% paraformaldehyde at 4 °C overnight and transferred to 30% sucrose for at least 16h at 4 °C. Organoids were then embedded in OCT, snap frozen in dry ice and ethanol, and stored at −80 °C. Frozen organoids were cryosectioned 15–20 µm thick, dried for at least 1 h and washed with PBS for OCT removal. Antigen retrieval was performed by steaming slices at 99 °C for 10 min in a 10 mM citrate solution at pH 6.0. Slides were washed and blocked in a 10% NDS, 0.3% Triton X-100 PBS solution and then incubated with primary antibodies at 4 °C overnight. Primary antibodies used in this study include: CTIP2 (Abcam, ab18465, Dilution 1:200); SOX2 (EMD Millipore, Dilution 1:200). The overnight incubation was followed by PBS washes and secondary antibody incubation at 1:200 in a 5% NDS, 0.15% Triton X-100 solution for 90 min at room temperature. Secondary antibodies used in this study include donkey anti-mouse Alexa Fluor 488 (Invitrogen) and donkey anti-rat Alexa Fluor 594 (Invitrogen, A21209, Lot 2078918). DAPI (2 µg/mL) was incubated for 20 min. Slides were mounted with Fluoromount-G (Thermo Scientific, 00-4958-02) and imaged on a Nikon W1 spinning disk confocal microscope or BZX710 Keyence epifluorescent microscope.

**Mass spectrometry sample preparation**

*Protein extraction and trypsin digestion.* Total protein was extracted from the organoids by sonication in lysis buffer (1% sodium dodecyl sulfate (SDS), 125 mM triethylammonium bicarbonate (TEAB), 75 mM NaCl, and HaltTM protease and phosphatase inhibitors). Lysates were cleared by centrifugation at 12,000 g for 15 min at 4 °C. Lysate samples from two additional control cell lines were included to improve protein detection. A different randomized block was used for each stage of sample processing, TMT labeling, and mass spec analysis, balanced for genotype and timepoint. Total protein was estimated using microBCA assay according to the manufacturer instructions (Cat. No. 23235, Thermo Fisher). 20 µg total protein from each sample was reduced with dithiothreitol (DTT), alkylated with iodoacetamide (IAA), and digested with trypsin on S-Traps (Protifi) per manufacture protocol.

*TMT labeling.* Samples were first assigned to one of five sets of TMTPro16 blocks (plex's), such that each plex was balanced for genotype and timepoint, with a random order in each plex (Supplementary Table 1). Two pooled reference controls were also included within each plex facilitating batch correction across plex's, except for plex 5 where four reference samples were included to complete the 16 channels. These reference controls were prepared by pooling equal amounts of lysate from all samples, which were digested separately, and then split in aliquots for TMT labeling. Peptide digests were reconstituted in 20 µl 100 mM TEAB and sonicated for 15 min at RT for TMT labeling. TMTPro reagents were brought to room temperature and dissolved in 20 µl Optima LC/MS-grade acetonitrile (ACN). 6 µl of ~59 mM TMTPro reagents was added to 20 µl reconstituted peptide accordingly. All TMT plex's were labeled at the same time. For the reference control, peptides were reconstituted in 80 µl. A total of 40 µl was labeled with 12 µl TMTPro 134 N. Labeling was performed at RT for 90 min with shaking at 400 rpm. The reaction was quenched by adding 3 µl 5% hydroxylamine/100 mM TEAB at RT for and shaken 15 min (6 µl for reference channel). All samples with in a plex were mixed along with 20 µg aliquots of each labeled pooled control, giving 320 µg total TMT labeled peptides per plex, and dried in a vacuum concentrator. Of this, 80 µg were used for peptide/protein quantification, and 240 µg was subject to phosphopeptide enrichment. Prior to peptide/protein quantification, the 80 µg aliquot was separated into 8 fraction using the Pierce High pH Reverse Phase Fractionation Kit per manufacturer protocol. The resulting fraction were dried down in a speed vac, and resuspended in 3% ACN, 0.1% formic acid.

*Phosphopeptides enrichment.* For phosphopeptide enrichment, the 240 µg TMT-labeled peptide aliquot were desalted with C18 spin column. Samples were resuspended and sonicated for 10 min in 300 µl buffer A (LC MS water 0.1% TFA) with 2% ACN. Resin was packed by centrifugation at 5,000 g for 2 min, activated twice with 300 µl ACN, and conditioned twice with 300 µl buffer A. Peptides were loaded once by centrifugation at 3,000 g for 1 min. Column was washed three times with 300 µl buffer A, and twice with 300 µl 5% methanol buffer A. Peptides were eluted twice with 300 µl 50% ACN 0.1% TFA and dried in a vacuum concentrator. Phosphopeptides were enriched using the AssayMAP Agilent Bravo instrument with Fe(III)-NTA IMAC resin according to the manufacturer's instructions (Agilent, Cat. No., G5496-60085). Peptides were resuspended in 210 µl 80% ACN 0.1% TFA, bath sonicated for 10 minutes, vortexed for 10 minutes, and quickly centrifuged. The resin was primed with 50% ACN 0.1% TFA and equilibrated with equilibration/wash buffer 80% ACN 0.1% TFA. Samples were loaded into the syringes then dispensed into the Flow Through Collection plate. Syringes were washed with the Cartridge Wash Buffer discarded into waste, then repeated with Syringe Wash Buffer. Elution buffer was aspirated into the syringes and dispensed into the Eluate Collection plate. Phosphopeptides were eluted in 18 µl 1% ammonium hydroxide solution and acidified with 1.8 µl 20% FA, and subject to high pH reverse phase fractionation as described above but condensed to 5 fractions.

Supplementary figures

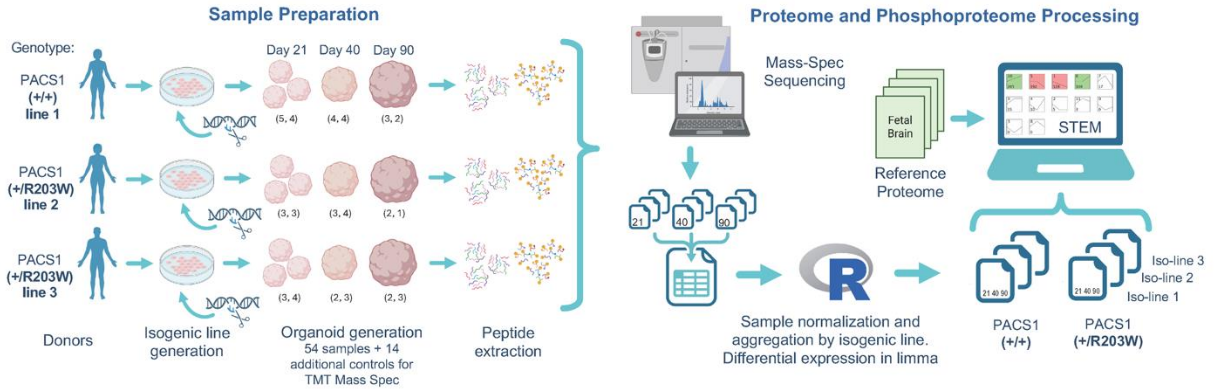


**Figure S1. Experimental design summary for sample preparation and proteome analysis.** Organoids were generated from three pairs of isogenic lines (PACS1$^{(+/+)}$ and PACS1$^{(+/R203W)}$) iPSC lines were obtained from one neurotypical and two p.R203W patient donor and CRISPR-edited to either insert or correct the p.R203W pathogenic variant. Organoids were collected on days 21, 40, and 90. Multiple organoids were pooled for day 21 samples due to the smaller organoid size. Numbers in parenthesis indicate the independent replicates obtained per timepoint, per PACS1$^{(+/+)}$ and PACS1$^{(+/R203W)}$ genotypes, yielding a total of 54 samples. Protein and phosphorylated peptides were processed through mass spectrometry. Outputs were normalized and aggregated by isogenic pair and differentiation in R using limma's linear model fit. Differential abundance tests were performed by timepoint. Normalized values were imported into the short time series (STEM) pipeline for time series expression trend clustering. Public data available for fetal brain development was also processed in R, analyzed on STEM and compared to the organoid proteome profiles. A similar criterion for grouping fetal samples based on region of origin and age was applied to homogenize the data.

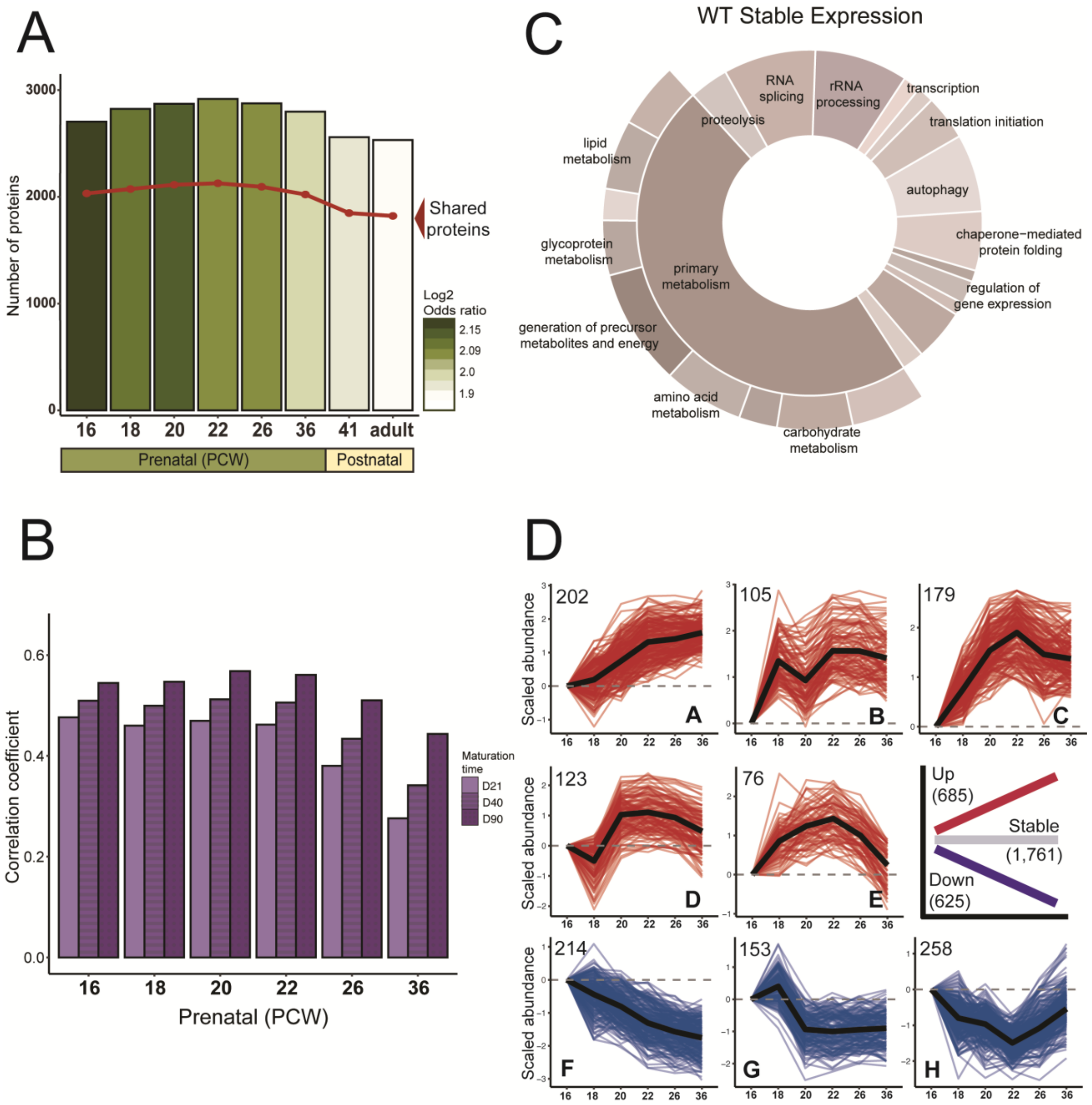


**Figure S2. Cortical organoids recapitulate proteome dynamics during brain development.** A) Total overlapping proteins between brain organoids and the reference proteome of the developing human brain. The bars represent the total number of proteins reported in the fetal brain by post-conception week (PWC), while the red line depicts the total number or protein identities shared with brain organoids. The color of the bar depicts the overlapping odds ratio between groups. B) Average Pearson's correlation coefficient of protein abundance between brain organoids and fetal brain developmental stages. C) Top GO terms in the stable cluster over maturation time in control organoids. D) Temporal clustering of expressed proteins in the fetal brain, up clusters A-E, down clusters F-H.

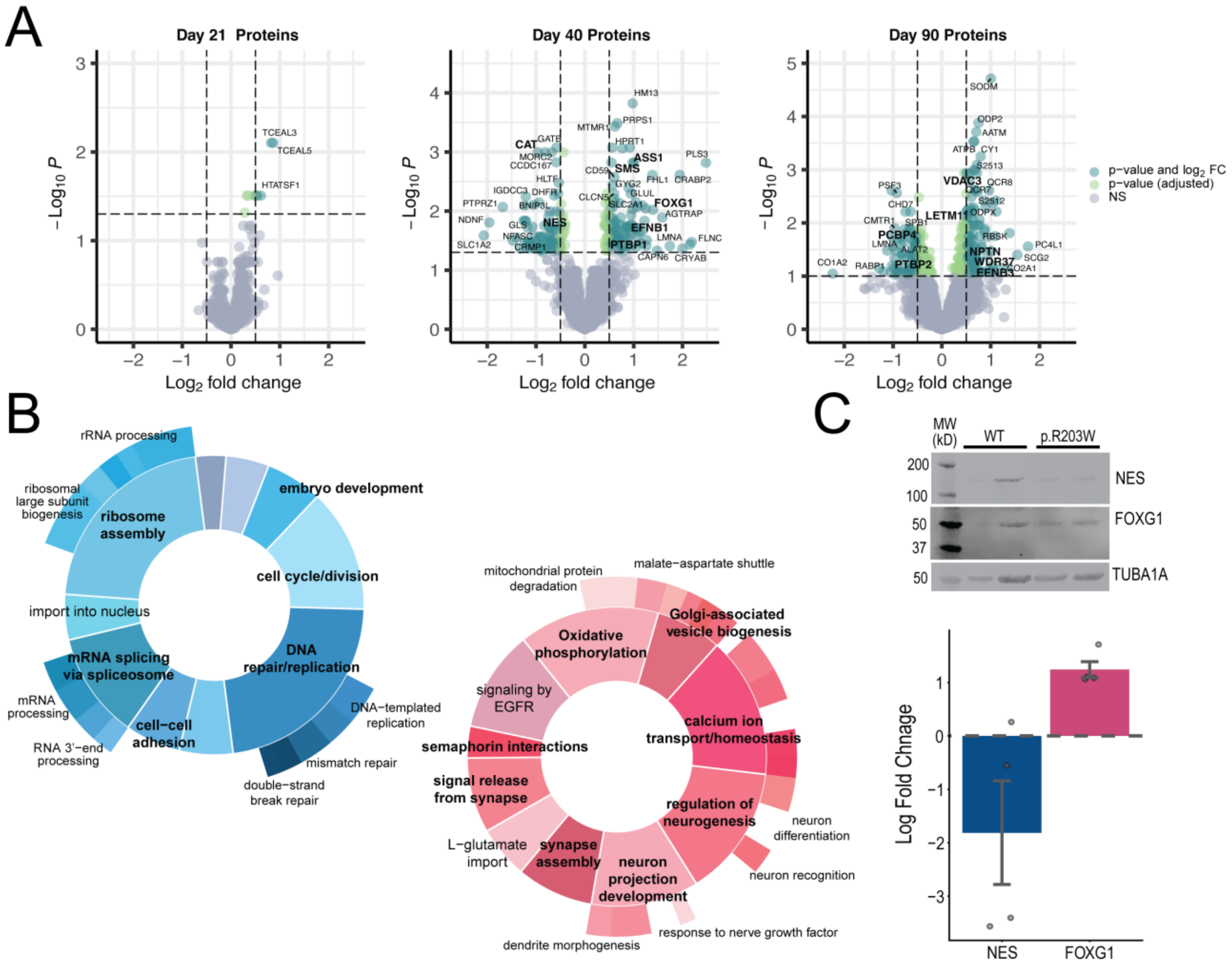


**Figure S3. Differentially expressed proteins in the p.R203W organoids during maturation.** A) Volcano plots of differentially expressed proteins in organoids carrying the p.R203W pathogenic variant compared to control organoids at day 21 (left volcano plot), day 40 and 90. (middle and right volcano plots). B) Summarized enrichment of biological processes that were downregulated (blue circle, upper left) and upregulated (red circle, lower right). C) Representative quantification of two differentially expressed proteins by Western Blotting in day 40 organoids (NES: downregulated, FOXG1: upregulated). Quantification of the signal was first normalized to $\alpha$-tubulin and then calculated the log2 fold change in the p.R203W pathogenic variant over the control.

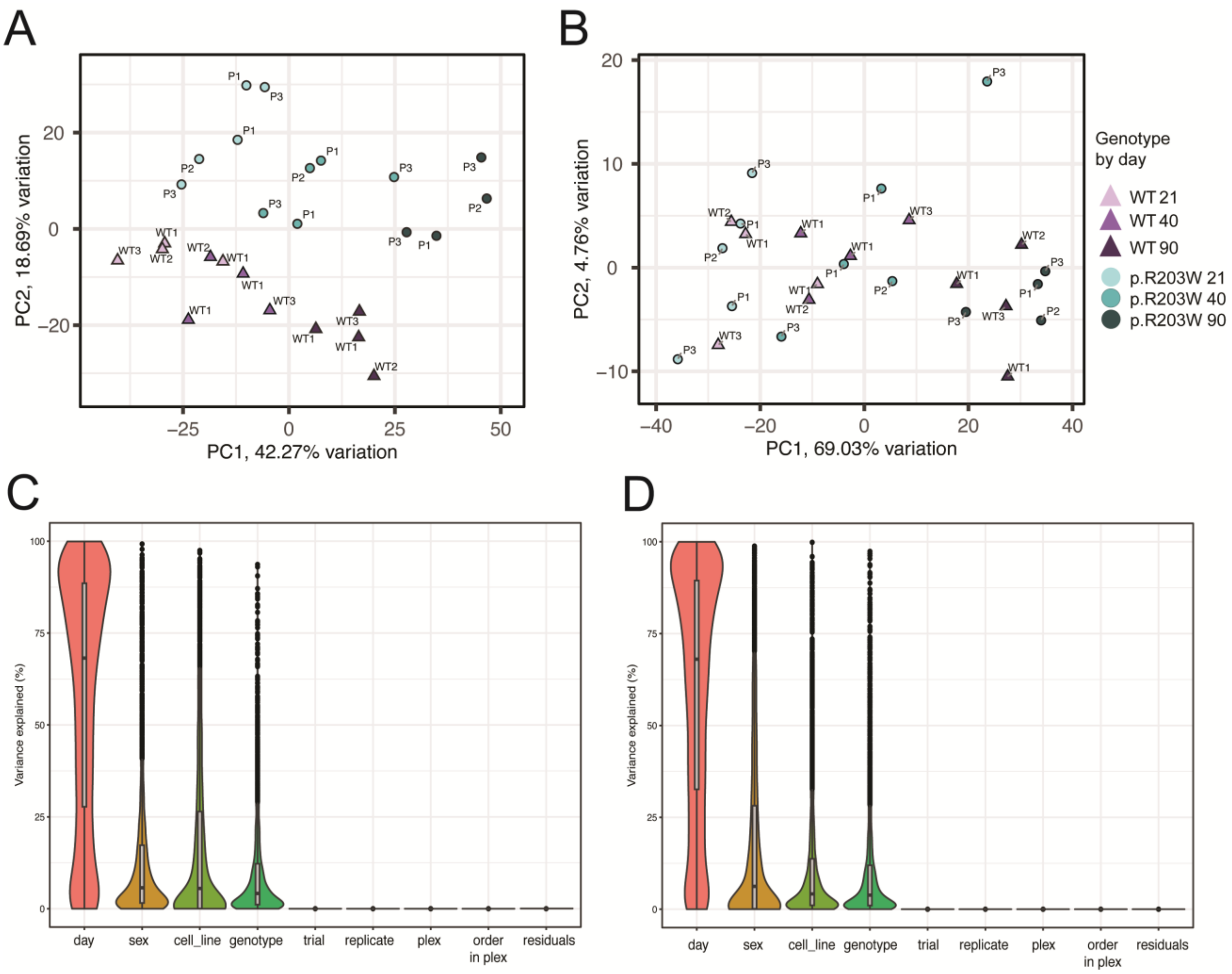


**Figure S4. Principal Component Analysis (PCA) of the aggregated organoid samples by isogenic line and differentiation and variance contribution by variable**. A) Proteins samples from WT and p.R203W are grouped by maturation time (component 1, ~42% of variation) and genotype (component 2, ~19% of variation). B) Phosphorylated peptides in all WT and p.R203W organoid samples, grouped by maturation time (component 1, ~69% of variation), while component 2 didn't have a strong association with either genotype or cell line (~5% of variation). Visualization of contribution to variability explained in each experimental variable after removal of technical batch effects using the limma:voom pipeline in the proteome (C) and phosphoproteome (D) datasets.

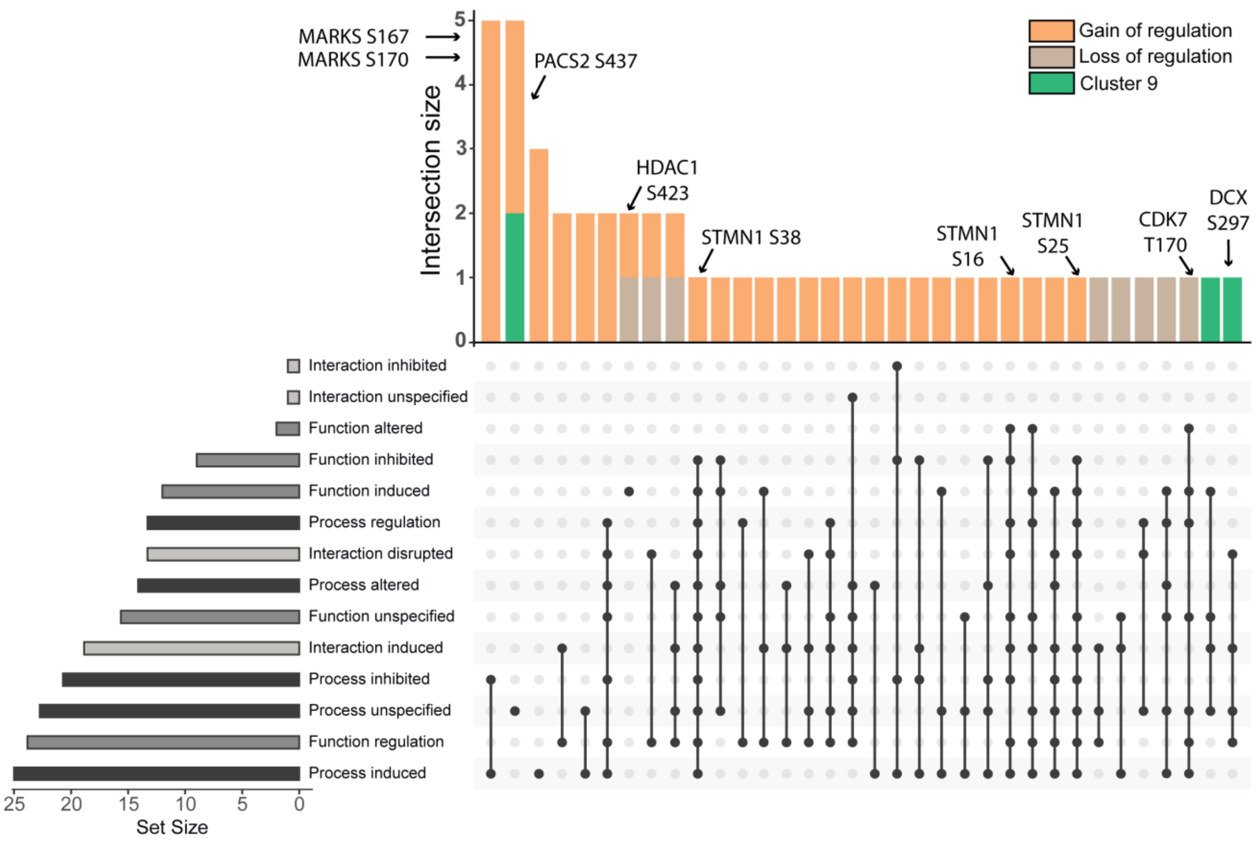


**Figure S5. Upset plot showing the relationship and distribution of reported processes, functions, and interactions of the phosphorylation sites with annotations reported in Phosphosite Plus database.** The top barplot (intersection size) indicates the total number of phosphorylation sites that share a given activity category represented in the bottom barplot (frequency of an annotation). The dark circles indicate all the shared categories among a given number of phosphorylated peptides. Some representative phosphorylated peptides are shown.

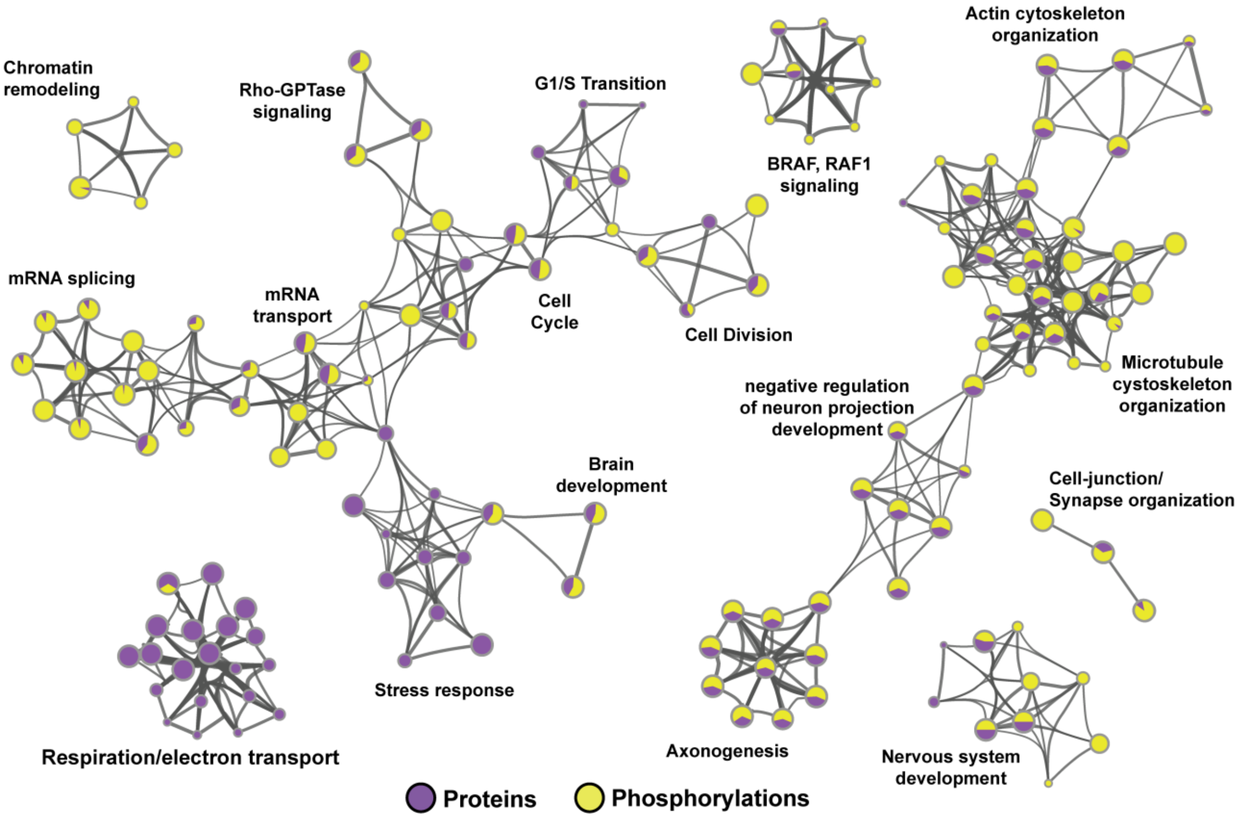


**Figure S6. Proteins and phosphorylation sites dysregulated by PACS1 p.R203W in organoids relate to convergent biological processes.** GO-term network showing the relationship between the top dysregulated processes in the organoids carrying the PACS1 p.R203W variant. Each circle represents a unique GO-term, and the edges represent the level of connectedness between terms. The size of the circle is associated to the gene list size. The color of each circle represents the proportion of representation in the proteome (purple) and phosphoproteome (yellow).
